# Inhibitor of apoptosis, *IAP*, genes play a critical role in the survival of HIV-infected macrophages

**DOI:** 10.1101/543017

**Authors:** Ramon Edwin Caballero, Simon Dong, Niranjala Gajanayaka, Hamza Ali, Edana Cassol, William D Cameron, Robert Korneluk, Michel J. Tremblay, Jonathan Angel, Ashok Kumar

**Affiliations:** Department of Biochemistry, Microbiology, and Immunology, Faculty of Medicine, University of Ottawa; Apoptosis Research Centre, Children’s Hospital of Eastern Ontario Research Institute, Ottawa; Department of Health Sciences, Carleton University, Ottawa, Canada; Division of Infectious Diseases, The Ottawa Hospital, and the Ottawa Hospital Research Institute, Ottawa, Ontario, Canada; Axe des Maladies Infectieuses et Immunitaires, Centre de Recherche du Centre Hospitalier Universitaire de Québec-Université Laval, and Département de microbiologie et immunologie, Faculté de médecine, Université Laval, Québec, Canada; Department of Pathology and Laboratory Medicine, Faculty of Medicine, University of Ottawa, Ottawa, ON, Canada

## Abstract

Latent viral reservoirs of HIV-1 that persist despite antiretroviral therapy (ART) are major barriers for a successful cure. Macrophages serve as viral reservoirs due to their resistance to apoptosis and HIV-cytopathic effects. We have previously shown that inhibitor of apoptosis proteins (IAPs) confer resistance to HIV-Vpr-induced apoptosis in normal macrophages. Herein, we show that second mitochondrial activator of caspases (SMAC)-mimetics (SM) specifically induce apoptosis of monocyte-derived macrophages (MDMs) infected *in vitro* with a R5-tropic laboratory strain expressing heat stable antigen, and GFP-expressing HIV, chronically infected U1 cells, and *ex-vivo* derived MDMs from naïve and ART-treated HIV patients. SM-induced cell death was found to be mediated by IAPs using IAP siRNAs, was independent of endogenously produced TNFα and was attributed to the concomitant downregulation of IAP-1/2 and the receptor interacting protein kinase-1 degradation following HIV infection. Altogether, modulation of the IAP pathways may be a potential strategy for selective killing of HIV-infected macrophages *in vivo*.

**Summary:** After more than 30 years of rigorous and intensive research since the identification of HIV-1, much progress has been made in understanding and controlling the pathogenesis of the virus. However, successful cure is currently unavailable. HIV-1 can remain undetected in various cell types, including memory T cells and macrophages, which make it difficult to achieve viral clearance without inciting cell death in infected cells. The “shock and kill” approach aims to reawaken dormant integrated virus and boosts host’s immune system for viral clearance in latently infected CD4+ T cells. However, to completely eradicate HIV in infected individuals, it is imperative to eliminate both CD4+ T cells and myeloid tissue reservoirs. Here we show that inhibition of the inhibitor of apoptosis (IAP) pathway, a cellular signalling pathway responsible for controlling cell death, by IAP inhibitors, smac mimetics can be utilized to kill HIV-infected macrophages. Deletion of cellular IAP proteins using smac mimetic, a synthetic anti-cancer compound currently being tested in several clinical trials, rendered HIV-infected macrophages susceptible to cell death. Herein, our results suggest that modulation of the IAP-associated signaling pathways may be a potential strategy for selective killing of HIV-infected macrophages.

## Introduction

Macrophages (Mϕ) are permissive to productive infection with HIV and a source of viral progeny for transmission to other cell types such as T cells [1–7]. HIV-infected Mϕ are widely distributed in tissues such as gastrointestinal and other mucosal tissues, lymph nodes and within the central nervous system where they have a life span extending from months to years [8–13]. In contrast to the characteristic depletion of CD4+ T cells, Mϕ do not decline in number, are resistant to apoptosis, survive active viral replication, and harbor unintegrated and integrated viral DNA in a state of latency [1,2,14–23]. In patients on effective antiretroviral therapy (ART), Mϕ serve as reservoirs as HIV persists in these cells, shielded against various host anti-viral responses and respond poorly to ART [1–3,15,24]. Moreover, infected Mϕ accumulate and retain virions within unique compartments designated as virus-containing compartments (VCCs) [25,26]. The virions present in VCC are protected from neutralizing antibodies and are inaccessible to anti-viral drugs [27–29]. Since HIV-infected Mϕ are not cleared by CD8^+^ T cells, neither current ART nor the immune system is able to effectively eliminate this reservoir [24].

While several recent studies support that Mϕ serve as a major non-T cell HIV reservoir [30–38], the role of Mϕ in HIV infection and persistence has been conclusively demonstrated by employing humanized BLT and myeloid only mice (MoM mice containing myeloid cells devoid of T cells). Honeycutt et al show that replication competent virus could be recovered from tissue Mϕ, and the transfer of infected Mϕ into uninfected animals resulted in sustained infection demonstrating that Mϕ are genuine targets for HIV infection *in vivo* [3]. Further, they demonstrated that HIV persists in Mϕ following suppressive ART *in vivo* in MoM model [39]. Therefore, to completely eradicate HIV in individuals on ART, it is imperative to eliminate both CD4+ T cells and myeloid tissue reservoirs. Most research to date has focused on eliminating the latent reservoir of CD4+ T cells by employing strategies to reactivate HIV in T cells and elimination of reactivated HIV-infected cells by host immunity [40–42]. However, approaches towards killing of HIV-infected Mϕ *in vitro* or *in vivo* are not well studied. Two recent studies have attempted to clear Mϕ reservoir by targeting infected Mϕ with CSF-1 receptor antagonists [43] and galactin-3 [44] with some success.

In order to devise strategies to eliminate HIV-infected Mϕ, it is imperative to identify apoptosis-related genes and signaling proteins involved in resistance of HIV-infected Mϕ to apoptosis. The mechanism underlying resistance of infected Mϕ to HIV-induced apoptosis may relate to the differential expression of pro-and anti-apoptotic genes including inhibitors of apoptosis (IAP) proteins [15,45]. The role of IAPs has been studied by employing antagonists of second mitochondria-derived activator of caspases (Smac), Smac mimetics (SM). SMs are small peptides that competitively inhibit Smac-IAP-1/2 interactions and repress anti-apoptotic functions of IAP proteins. Recently, IAP1/2 and survivin, another member of the IAP family were suggested to be involved in survival of HIV-infected CD4+ T cells [46,47]. In addition, IAPs have been implicated in protection against hepatitis B infection and in the reversal of HIV latency in CD4+ T cells [48,49]. Using HIV-Vpr as an apoptosis-inducing agent, we have shown a protective role for IAP genes in resistance to cell death in Mϕ [50–52]. CpG-induced protection against apoptosis and mitochondrial depolarization in monocytic cells was shown to be mediated by c-IAP-2 induction [50,52]. Moreover, down regulation of IAP-1/2, by using siRNAs and SMs, sensitized Mϕ to Vpr-induced apoptosis [51]. Therefore, strategies based on suppressing IAPs by employing SMs, may be useful in killing HIV-infected Mϕ. Herein, we show that SMs induced apoptosis in *in vitro* HIV-infected Mϕ and that this may occur through the concomitant down regulation of both IAPs and receptor interacting protein kinase-1 (RIPK-1).

## Results

### SMs induce cell death in HIV-infected myeloid U1 cells but not in counterpart uninfected U937 cells

SMs bind to cIAP1/2 and promote their E3 ligase activity which leads to their auto-ubiquitination, subsequent proteasome degradation and apoptosis [53,54]. We have previously shown that cIAP1/2 genes play a protective role in mediating survival of Mϕ in response to Vpr-induced cell death [50–52]. To determine whether SMs impact apoptosis in HIV-infected Mϕ, chronically infected U1 cells and uninfected counterpart U937 cells were treated with SM-LCL161 followed by assessment of cell death by PI staining and flow cytometry. SM treatment induced significant cell death in U1 cells but not in U937 cells (Fig 1A). To determine whether differentiation of U937 render these cells susceptible to SM-induced apoptosis, U937 and U1 cells were differentiated with PMA. Similar to the effect of SM on undifferentiated U1 cells, SM-LCL161 induced significant cell death in differentiated U1 cells but not in differentiated U937 cells (Fig 1B). Specific killing of HIV-infected U1 cells was further confirmed by showing cleavage of caspase-3 in U1 but not in U937 cells (Fig 1C).

**Figure 1.**
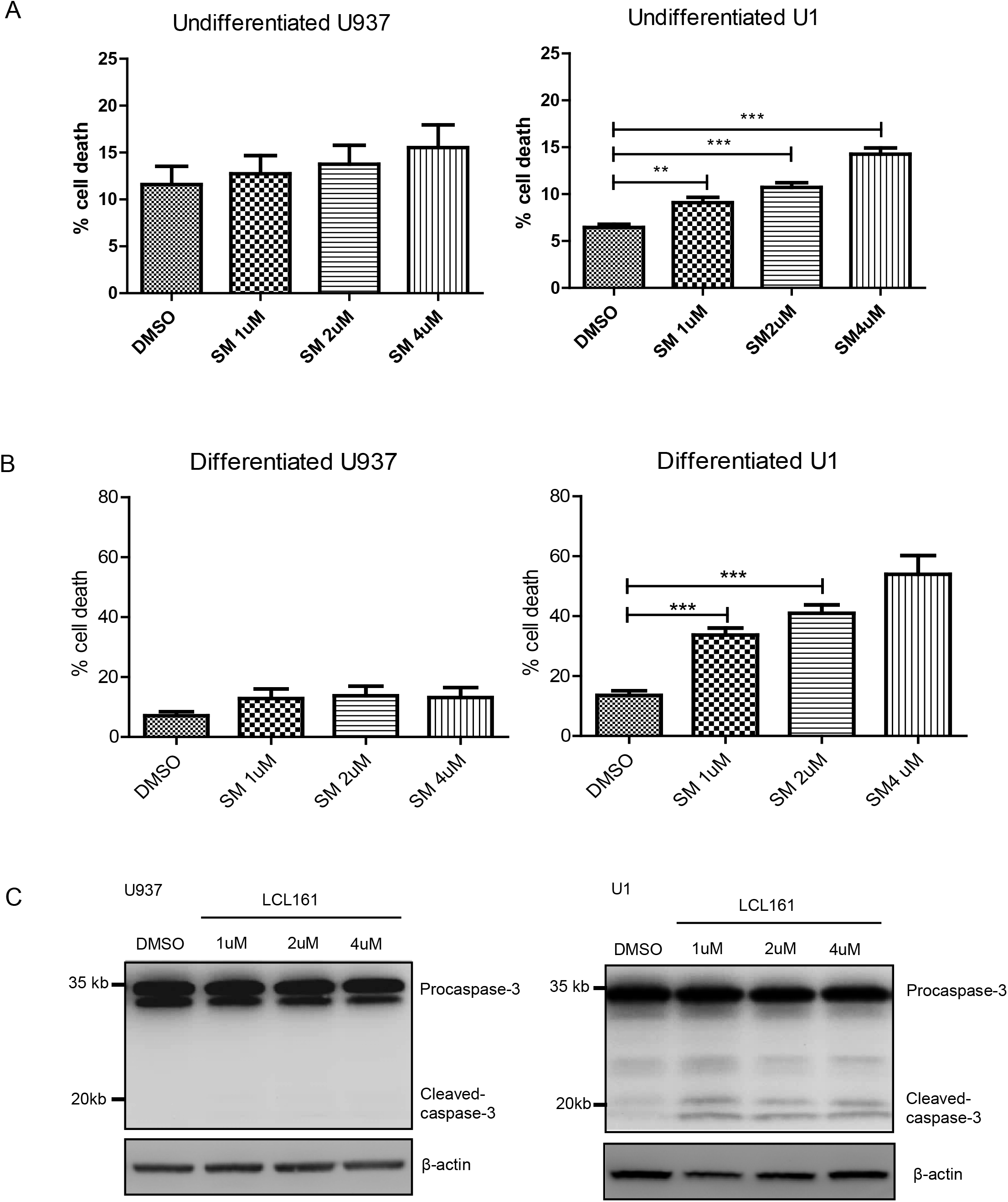
SM induces cell death of HIV-infected myeloid cells. (A) U937 (n=9) and chronically infected counterpart U1 cells (n=10) were treated with SM-LCL161 at 1, 2, and 4 μM for 48 hr. (B) PMA differentiated U937 (n=7) and U1 cells (n=11) were treated with SM LCL161 at 1, 2, and 4 μM for 48 hr. Cell death was assessed by intracellular PI staining. The p-values were calculated using Mann-Whitney U test. (***p=<0.0001, **p=0.002). (C) U937 and U1 were treated with increasing concentration of SM-LCL161 for 48 hr and cytosolic fractions were collected and subjected to Western immunoblotting. 30 µg of total proteins were loaded to the protein gels. The membranes were probed with antibodies specific for caspase-3. The figure shown is a representative of three experiments.

### SMs induce cell death in ***in vitro*** HIV-infected MDMs and MDMs derived from HIV-infected patients

To validate above findings in primary MDMs, we first verified the functional activity of SM by treating HIV-infected MDMs with LCL161 and observed degradation of both cIAP1 and cIAP2 (Fig 2A) as reported earlier [51,55]. The *in-vitro* HIV_CS204-_infected MDMs were treated with SM-LCL161 followed by assessment of cell death by PI staining and flow cytometry. SM-LCL161 induced significant cell death of HIV_CS204_-infected MDMs but not in mock-infected MDMs (Fig 2B). Representative histograms of the intracellular PI staining are shown (Fig. 2C). The p24 values in MDMs infected with HIV_CS204_ for 7 days are shown in Fig 2D. Further, to determine whether Mϕ derived from HIV-infected individuals are similarly prone to SM-induced cell death, MDMs were generated from ART treated and ART naïve HIV-infected individuals and treated with SM-LCL161. Consistent with *in vitro* infection studies, *ex vivo* derived MDMs from treatment naïve and ART-treated HIV-infected individuals showed significantly increased susceptibility to SM-LCL161-induced cell death in a dose-dependent manner (Fig 2E).

**Figure 2.**
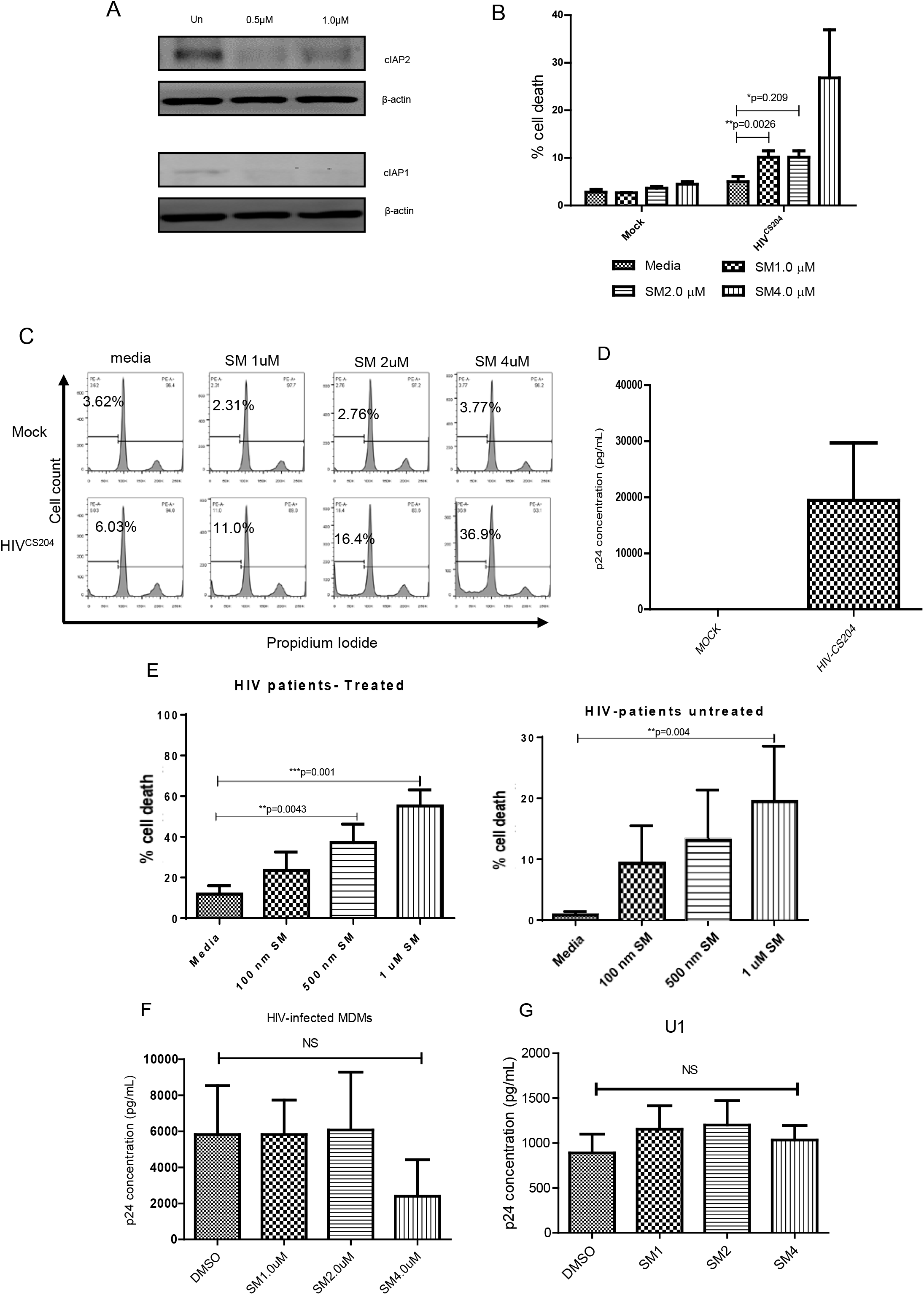
SM induces cell death of HIV-infected MDMs. (A). The HIV-infected MDMs were treated with increasing concentration of SM-LCL161 for 48 hr. The cytosolic fractions were subjected to Western immunoblotting. The membranes were probed with antibodies specific for human cIAP-1 and cIAP-2. (B) Human MDMs were *in vitro* infected with HIV-_CS204_ (100 ng p24 / well) for 7 days. The cells were then treated with SM-LCL161 for 48 hr and cell death was assessed by PI staining and flow cytometry. The representative histograms are shown in (C). (D) After 7-days of infection, supernatants were analyzed for p24 by ELISA. (E) MDMs generated from naïve and ART-treated HIV-individuals were treated with SM-LCL161 for 48 hr. Cell death was assessed by PI staining and flow cytometry. Supernatants from SM-treated *in vitro* HIV-infected MDMs (F) and SM treated U1 cells (G) were analyzed for p24 secretion. The p-values were calculated using Mann-Whitney U test.

Apoptosis has been shown to induce viral activation and replication in latently infected U1 and ACH2 cell lines [56]. In addition, Pache et al have shown that SMs can affect viral transcription in infected CD4^+^ T cells via NF-κB dependent signalling [49]. To determine if SMs affect HIV replication in Mϕ, *in vitro* HIV-infected MDM were treated with SM -LCL161 for 48 hr followed by analysis of p24 secretion. Interestingly, virus replication in primary HIV-infected MDM (Fig 2F) and in HIV-infected U1 cells (Fig 2G) was not affected by SM treatment.

### Smac mimetics specifically kill HIV-infected MDMs

Based on above results, it is unclear if SMs are killing HIV-infected and/or bystander uninfected HIV-exposed MDM. To examine this, we employed a R5 laboratory strain of HIV-1, HIV-Bal-HSA, expressing mouse HSA (CD24). Expression of CD24 by HIV-infected cells can be used to identify infected cells by flow cytometry using FITC-conjugated anti-mouse HSA antibody [57]. MDMs were infected with HIV-Bal-HSA for 7 days followed by treatment with SM-AEG 40730 for another two days. Specific killing of HSA-expressing (HIV-infected) cells by SM-AEG 40730 was quantified by counter staining with Annexin-V labelled with BV-711. To rule out non-specific low level fluorescence by dead cells/debris, highly intense FITC positive HSA-expressing cells were gated and further quantified for Annexin-V expression. The gating strategy is shown in Fig. 3A. Quantification of Annexin-V positive, total (HSA+ and HSA-; Fig 3B, left panel) and intensely HSA+ MDMs (Fig 3C, left panel) revealed that SM-AEG 40730 killed significantly higher number of total HIV-infected (HSA+ and HSA-) and high HSA-expressing MDMs compared to the DMSO-treated HIV-infected cells. Representative histogram showing killing of total HSA+ and HSA-(Fig 3B, right panel) and intensely positive (Fig 3C, right panel) HSA-expressing cells is shown.

**Figure 3.**
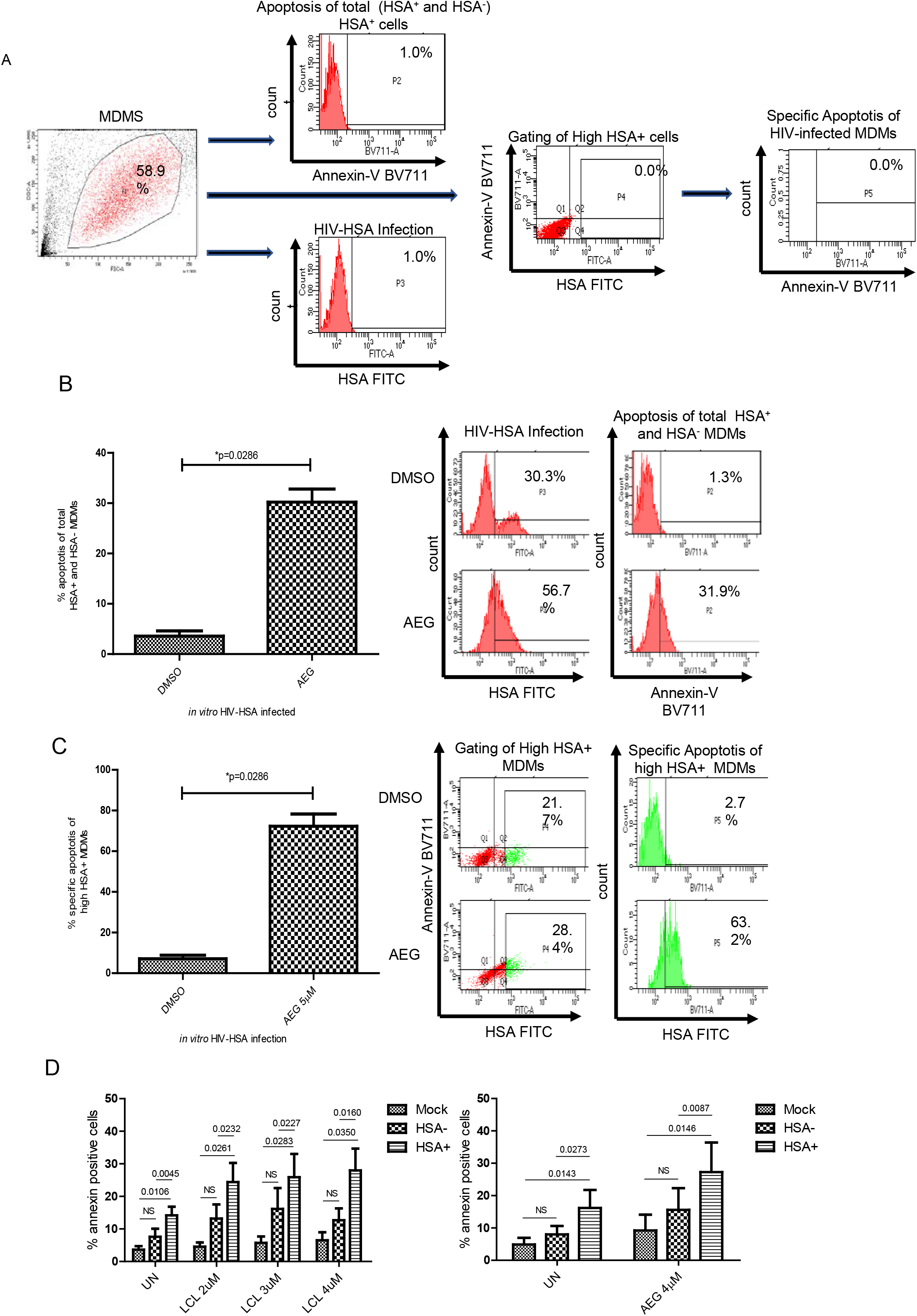
SM specifically induces cell death of HIV-HSA-infected MDMs. (A) Gating strategy for the detection of apoptosis of HIV infected total (HSA+ and HSA-) MDMs, and high HSA-expressing MDMs. (B) MDMs were *in vitro* infected with HIV-Bal-HSA for 9 days. Cells were treated with DMSO or 5µM AEG40730 for 72 hr. Cell death of total HIV-HSA+ and HIV-HSA-MDMs was detected by Annexin-V-BV711 and flow cytometry (left panel). Representative histogram shows HIV-HSA-infected cells (middle panel) and cell death of total HSA+ and HSA-HIV-infected cells (right panel). (C) Intensely HSA positive HIV-infected cells were gated and analyzed for apoptosis by Annexin-V-BV711 staining and flow cytometry (left panel). Representative histogram shows intensely positive HIV-HSA-infected cells (middle panel) and cell death of high HSA+ cells (right panel). The p values in B and C were calculated using Mann-Whitney U test. (D) SM specifically kill HIV-HSA-infected cells but not HIV-HSA-MDM. MDMs were infected with HIV-HAS for 11 days followed by treatment with either SM-LCL-161 (upper panel) or AEG40730 (lower panel) for another two hr followed by analysis of cell death by PI staining (n=5). The p-values were calculated using paired t test.

To determine whether SMs killed uninfected HIV-exposed bystander cells, MDM were infected with HIV-Bal-HSA for 7 days followed by treatment with either SM-AEG40730 or SM-LCL161 for another two days. Specific killing of HSA-expressing (ie HIV-infected) and HSA-negative (HIV uninfected) cells by SM-AEG 40730 or SM-LCL161 was quantified by counter staining with BV-711 labelled Annexin-V as above. SM-AEG40730 and SM-LCL161 killed significantly high numbers of HIV-HSA-expressing (HIV-infected) cells compared to either the mock or HSA-negative (HIV-uninfected/bystander, HIV-exposed) cells (Fig 3 D). However, killing of HSA-negative (HIV-uninfected/bystander) cells was relatively higher than mock-infected cells but was not significant suggesting that SMs specifically kill HIV-infected Mϕ.

Similar experiments were performed with a GFP-expressing HIV strain, HIV-eGFP. MDMs were infected with HIV-eGFP for 7 days followed by treatment with SM-AEG 40730 for two days. Expression of GFP by HIV-infected cells can be visualized by flow cytometry to identify infected cells. Specific killing of GFP-expressing (HIV-infected) cells by SM was quantified by counter staining with BV-711 labelled Annexin-V. Highly intense GFP positive cells were gated and further quantified for Annexin-V expression. The gating strategy is shown in Fig 4A. MDMs infected with HIV-GFP for 1-2 days could be detected by flow cytometry; however, MDMs infected for 7 days could not be detected as the virus multiplied for one round only (data not shown). Interestingly, treatment of HIV-GFP-infected cells with SM-AEG 40730 revealed killing of significantly high number of total HIV-GFP+ and GFP- MDMs compared to the DMSO-treated HIV-GFP-infected MDMs (Fig 4B, left panel). Specific killing of HIV-GFP infected cells by SM-AEG 40730 by gating dual GFP-positive and Annexin-V+ MDMs revealed that SM-AEG 40730 killed significantly higher number of intensely GFP+ HIV-infected cells compared to the DMSO-treated HIV-infected cells (Fig 4C, left panel). Representative histogram showing killing of total GFP+ and GFP-MDMs (Fig 4B right panel) and intensely GFP+, HIV-GFP-infected cells (Fig 4C, right panel) by SM-AEG40730 is shown.

**Figure 4.**
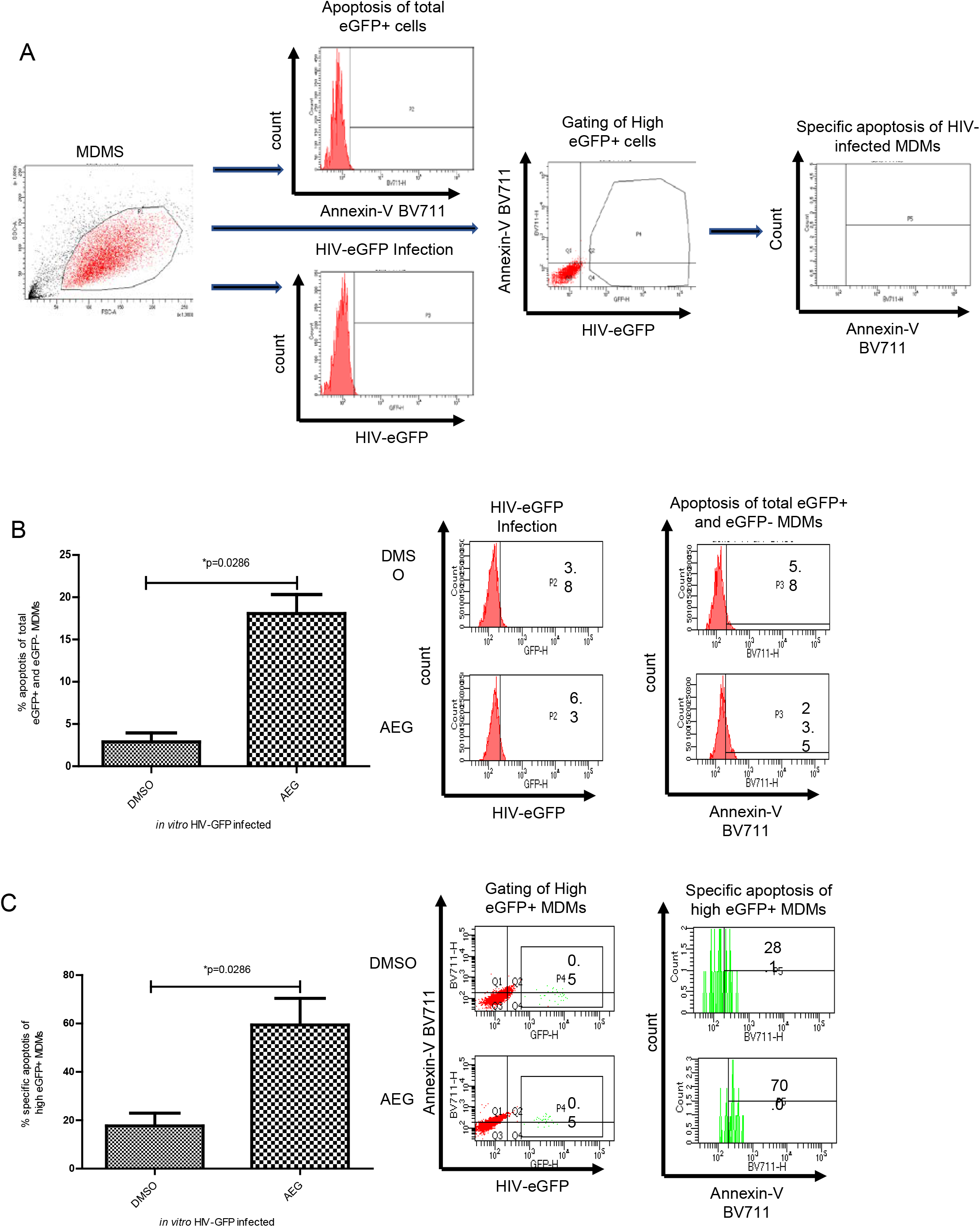
SM specifically induces cell death of HIV-GFP-infected MDMs. (A) Gating strategy for the detection of apoptosis of total HIV-eGFP infected and specific high HIV-GFP expressing MDMs. (B) MDMs were *in vitro* infected with HIV-eGFP for 7 days. Cells were treated with DMSO or AEG40730 for 72 hr. The cell death of the total population of HIV-GFP-infected MDMs was detected by Annexin-V-BV711 and flowcytometry (left panel). Representative histogram shows HIV-GFP-infected cells (middle panel) and cell death of total GFP+ and GFP-HIV-infected cells (right panel). (C) Intensely GFP-positive HIV-infected cells were gated and analyzed for apoptosis by Annexin-V-BV711 staining and flow cytometry (left panel). Representative histogram showing intensely positive HIV-GFP-infected cells (middle panel) and cell death of high GFP+ cells (right panel). p-values were calculated using Mann Whitney U test (n=4).

### Knocking down IAP genes results in specific killing of HIV-infected MDM

To confirm the involvement of IAPs in SM-mediated killing of HIV-infected MDMs, we employed IAP-1/2 siRNAs as shown previously [50–52]. MDMs generated from PBMC from healthy donors were infected with HIV-Bal-HSA for 7 days followed by treatment with either non-targeting siRNA or IAP siRNAs for 72 hrs. Killing of HIV-infected cells in the presence of IAP siRNA treated cells was analyzed by staining with Annexin-V labelled with BV-711 as above. MDMs intensely expressing HSA and Annexin-V were gated and quantified as described above. The gating strategy is shown in Fig 5A. Quantification of Annexin-V positive total HSA+ and HSA-and intensely HSA-positive MDMs revealed that knocking down IAPs by siRNAs killed significantly higher number of total HIV-HSA+ and HSA-(HIV infected) MDMs (Fig 5B, left panel) and intensely HSA+ MDMs (Fig 5C, left panel) compared to the control HIV-HSA-infected cells treated with non-targeting siRNAs. Representative histogram showing killing of total HSA+ and HSA-(Fig 5B, right panel) and intensely HSA positive (Fig 5C, right panel) MDMs is shown.

**Figure 5.**
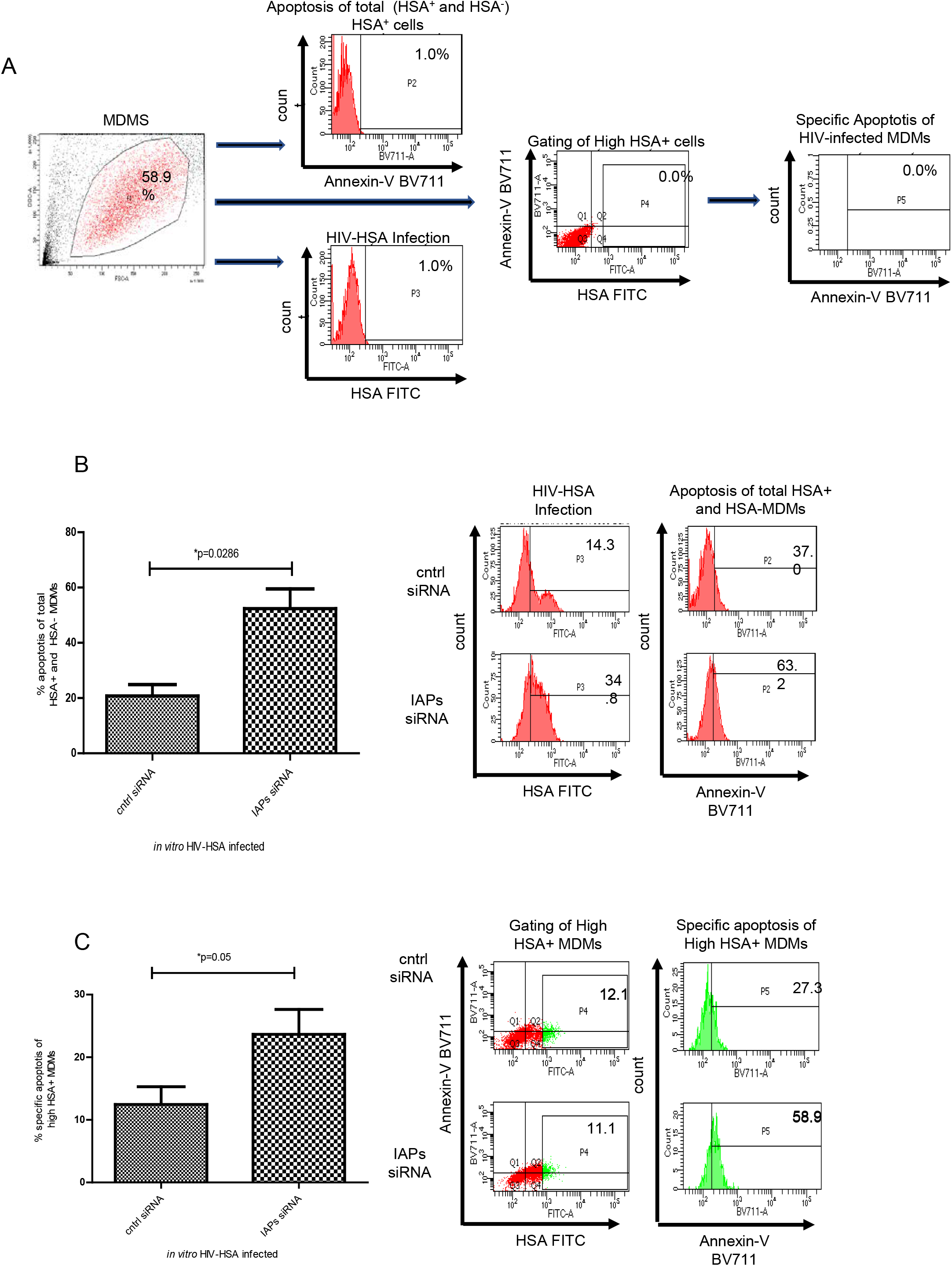
IAP siRNA specifically induces cell death of HIV-HSA-infected MDMs. (A) Gating strategy for the detection of apoptosis of total HIV-HSA infected MDMs, HSA-expressing MDMs, and specific apoptosis of HIV-HSA infected MDMs. (B) MDMs were in vitro infected with HIVNL4.3-Bal-HSA for 9 days. The cells were transfected with non-targeting control siRNA or cIAP1/2 siRNA. After 72 hr of transfection, cell death of the total population of HIV-HSA-infected MDMs was detected by Annexin-V-BV711 and flow cytometry (left panel). Representative histogram showing HIV-HSA-infected cells (middle panel) and cell death of total (HSA+ and HSA-) HIV-infected cells (right panel). (C) Intensely HIV-HSA positive HIV-infected cells were gated and analyzed for apoptosis by Annexin-V-BV711 staining and flow cytometry (left panel). Representative histogram showing intensely positive HIV-HSA-infected cells (middle panel) and cell death of high HSA+ cells (right panel). p-values were calculated using Mann Whitney U test (n=4).

### SM-induced cell death in HIV-infected MDM is mediated by apoptosis

To determine whether SM-induced cell death in *in vitro* HIV-infected MDM is due to apoptosis, caspase activation was quantified based on the fluorescent signal of cleaved caspase substrates. Treatment of HIV_cs204_-infected MDM with SM-LCL161 showed activation of caspases 3, 8, and 9 in contrast to the mock-infected MDM (Fig 6A-6C). Moreover, prior treatment with zVAD-FMK, a pan-caspase inhibitor, reduced the activation of caspase-8 and 9 after SM-LCL161 treatment (Fig 6B-C). A representative histogram for the induction of caspase 3, 8 and 9 following SM treatment of HIV-infected MDM is shown (supp. Fig 1).

**Figure 6.**
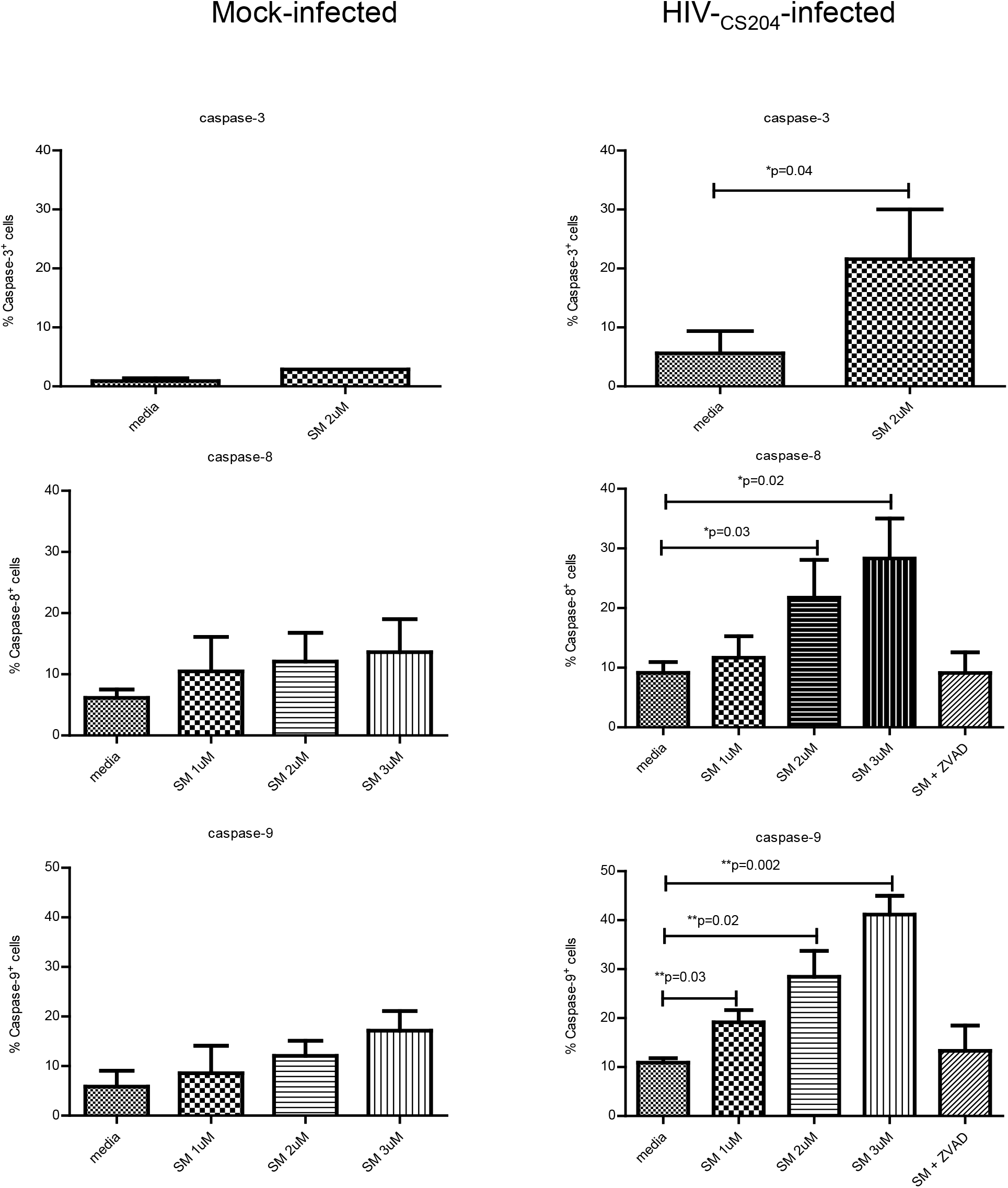
SM-induced cell death of HIV-infected MDMs is mediated by apoptosis. *In vitro* HIV-infected MDM were treated with SM-LCL161 for 48 hr. The cells were treated with Z-Vad pan-caspase inhibitor for 2 hr prior to SM-LCL-161 (3 μM). The activation of caspases was detected by fluorescent caspase substrate (caspase-3, n=3; caspase-8, n=4; caspase-9, n=6). The p values were calculated using Mann-Whitney U test.

### TNF-α mediates SM-induced apoptosis in U1 cells but not in primary HIV-infected MDM

SM-induced cell death of various tumor cells is mediated by endogenously produced TNF-□ following SM treatment through the activation of the non-canonical NF-κB pathway [58,59]. To determine if SM-induced apoptosis in HIV-infected MDM is due to endogenous TNFα production, SM-LCL161-treated U937, U1 cells and *in vitro* HIV-infected primary MDM were analyzed for TNF-α secretion. SM-LCL161 treatment resulted in low level although significant TNF-α production in undifferentiated and differentiated U937 and U1 cells (Fig 7A-D) in contrast to both *in vitro* mock-and HIV-infected MDM (Fig 7E). Similarly, *ex vivo* derived MDM from HIV-infected patients did not produce significantly higher levels of TNF-α following SM-LCL161 treatment compared to the untreated negative controls (Fig 7F).

**Figure 7.**
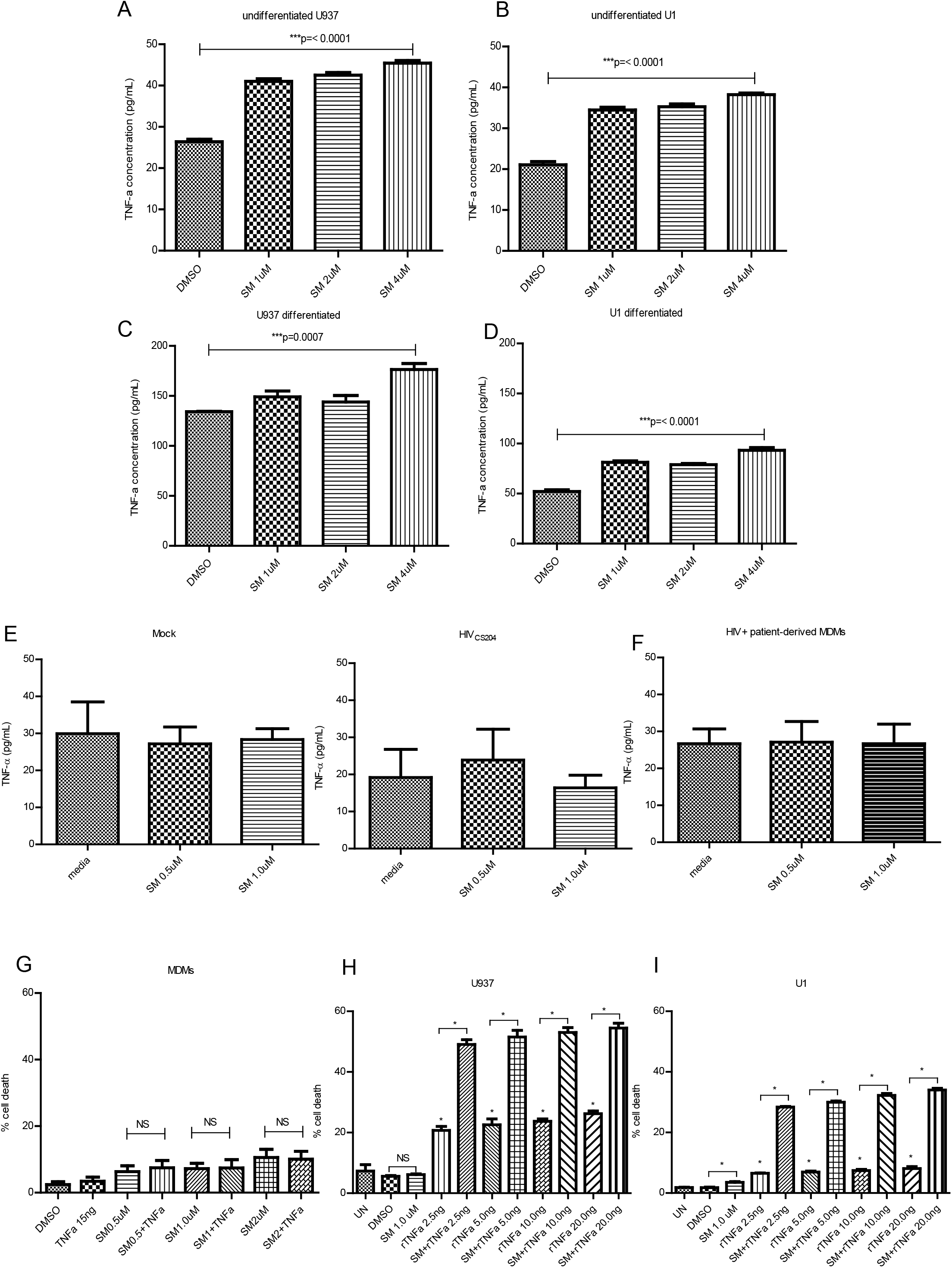
TNF-α mediates SM-induced apoptosis in U1 cells but not in primary HIV-infected MDM. SM induced cell death in U1 cells is regulated by r TNF-a. SM induce TNFα secretion in undifferentiated and differentiated U937 and U1 cells (A-D). Undifferentiated U 937 (A), undifferentiated U1 (B), differentiated U937 (C) and differentiated U1 (D) cells were treated with various concentrations of SM-LCL161 for 48 hr. The supernatants were analyzed for TNF-α production by ELISA. SM treatment of HIV-infected MDMs does not induce TNFα production. (E). Human MDMs were *in vitro* infected with HIV-_CS204_ (100 ng p24 / well) for 7 days followed by the addition of SM-LCL161 for 48 hr (n=3). (F) MDMs derived from HIV-patients were treated with SM-LCL161 for 48 hr (n=4). Supernatants were analyzed for TNFα production by ELISA. U 937 (G) and U1 (H) were treated with either SM-LCL161 alone, rTNF-α alone or with SM-LCL161 and various concentrations of rTNF-α for 48 hr followed by analysis of cell death by PI staining and flow cytometry. (I) MDMs were treated with SM-LCL161 for 2 hr followed by the addition of rTNF-α for 48 hr. Intracellular PI staining and flow cytometry were used to assessed levels of cell death (n=3). p-values were calculated using paired t test or Mann-Whitney U test.

To evaluate the impact of TNF-α in SM-induced apoptosis of primary MDM, SM-LCL161-treated MDM were stimulated with rTNF-α followed by analysis of cell death by PI staining. Treatment of MDMs with SM-LCL161 and TNFα did not result in cell death (Fig 7G). In contrast, rTNF-α either alone or in combination with SM-LCL161 induced significant cell death in U937 and U1 cells (Fig 7H, I) similar to that observed in various tumor cells [60,61]. These results suggest that SM-mediated killing of HIV-infected MDM is independent of TNFα.

### HIV-infected MDM do not develop M1 phenotype before or after SM treatment

Macrophages polarized with IFNγ develop a M1 phenotype which is highly susceptible to SM-induced cell death (Supp. Fig 2). Therefore, it is possible that SM-induced cell death of HIV-infected MDMs is due to the development of M1 phenotype following HIV infection. To determine whether HIV-infected MDM develop M1 phenotype before or after SM treatment, cytokine array analysis for the following cytokines was performed: IL-17F, GM-CSF, IFNγ, IL-10, CCL20/MIP3a, IL-12p70, IL-13, IL-15, IL-17a, IL-22, IL-9, IL-1β, IL-33, IL-21, IL-23, IL-5, IL-6, IL-17ε/IL-25, IL-27, IL-31, TNFα, TNFβ, and IL-28A. HIV-infected MDM secreted significantly high levels of CCL20/MIP3α, IL-6, and TNFα compared to the mock control. There was no significant difference in the secretion of IL-10, IL-21, IL-13, and IL-23 between the HIV-infected and mock-infected controls (Supp. Fig 3). Remaining cytokines were not detected in either group suggesting that HIV infection of MDMs does not result in the upregulation of cytokines related to M1 phenotype. SM treatment did not affect the secretion of above mentioned cytokines including CCL20/MIP3α, IL-6, IL-23, IL-10, IL-21, IL-13, and TNFα in *in-vitro* HIV-infected MDMs (Supp. Fig 4) or in *ex-vivo* derived MDMs from HIV-infected patients (Supp. Fig 5). These results suggest that *in-vitro* HIV-infected MDM either before or after SM treatment did not express M1 phenotype and SM-mediated apoptosis of HIV-infected MDM is independent of M1-polarization.

### HIV-infection downregulates RIPK1 in MDMs

SM-induced apoptosis of HIV-infected macrophages may be ascribed to the impaired expression of IAP-associated signalling kinases such as RIPK-1 [62,63]. RIPK-1 plays a key role in the regulation of various cellular processes such as NF-κB signalling and apoptosis [64]. Moreover RIPK-1 is a target substrate for HIV protease, a viral protein that is synthesized late in the viral life cycle and inactivates RIPK1 in HIV-infected primary CD4+ T cells [65]. To determine whether RIPK1 is similarly cleaved and inactivated in HIV-infected MDMs, *in vitro* mock and HIV_CS204_-infected MDMs for 7 days were treated with SM -LCL161 for 2 days followed by immunoprobing for RIPK-1. HIV infection resulted in the downregulation of RIPK-1 in the presence and absence of SM-LCL161 compared to the mock infected controls (Fig 8A). This was also demonstrated by *in vitro* infection of MDMs with HIV_CS204_ for 2-8 days. Infection with HIV_CS204_ resulted in cleavage of RIPK1 with a relative decrease in full length RIPK1 while the cleaved RIPK1product gradually increased over time (Fig 8B).

**Figure 8.**
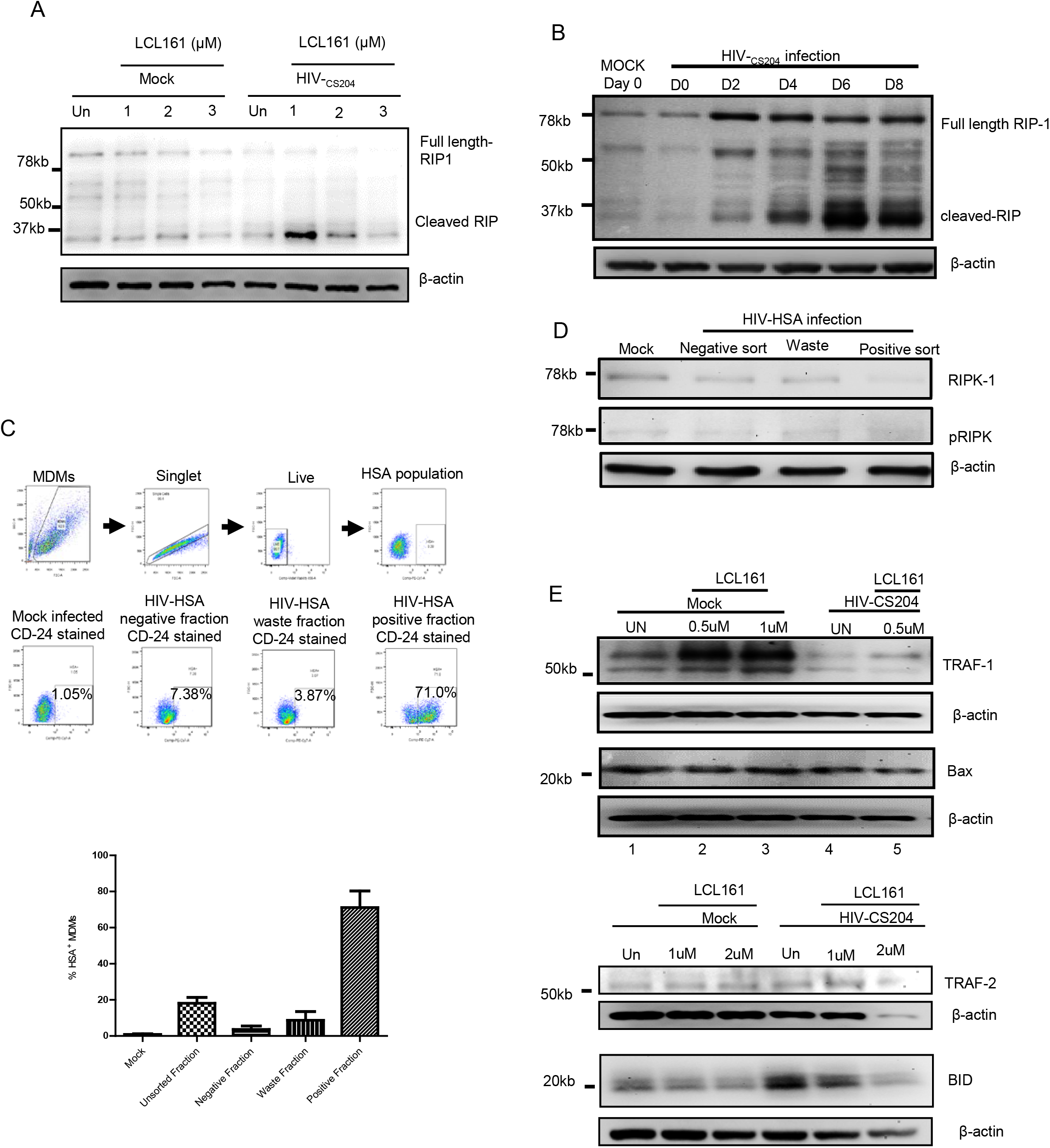
HIV infection results in downregulation of RIPK1 in MDMs. (A) Human MDMs were *in vitro* infected with HIV-_CS204_ (100 ng p24 / well) for 7 days followed by the addition of SM-LCL161 for 48 hr. (B) Human MDMs were *in vitro* infected with HIV-_CS204_ (100 ng p24 / well) and cells were harvested on day 0, 2, 4, 6, and 8. Cell lysates were subjected to Western immunoblotting for RIPK-1. The results shown are representative of 4 experiments (A) and twice for B. (C) Gating strategy for detection of HIV-HSA infected macrophages. HIV-infected bulk MDMs were gated as singlets followed by staining for live cells using E450 live/dead staining kit (Invitrogen). HIV-HSA-infected cells were detected within the live cell population by using FITC-labelled anti-CD24 antibodies (left panel). MDM were *in vitro* infected with R5 tropic HIV-Bal-HSA as above for 11 days following which cells were subjected to magnetic column separation using CD-24 (HSA)-biotin conjugated antibodies. The % of HIV-HSA-infected MDM in isolated unsorted, negative, waste and positive fractions as assessed by flow cytometry is shown (right panel). (D) Isolated fractions of HIV-HSA, namely negative fraction (negative sort), waste fraction and positively isolated fraction (positive sort) were subjected to Western immunoblotting for analysis of RIPK-1 and pRIPK-1. (E). Human MDMs were *in vitro* infected with HIV-_CS204_ (100 ng p24 / well) for 7 days followed by the addition of SM-LCL161 for 48 hr. Cell lysates were analyzed for TRAF-1, TRAF-2, Bid and Bax by Western immunoblotting. The results shown are a representative of two (upper panel) and 4 (lower panel) experiments respectively.

To confirm the downregulation of RIPK-1 in HIV-infected MDM, MDMs were infected with HIV-Bal-HSA. After 9 days of infection, HIV-infected HSA-expressing MDMs were harvested by magnetic column separation based on HSA expression followed by immunoblotting for RIPK-1 analysis [57]. The negative fraction represents HIV-exposed uninfected cells that do not express HSA on their surface, and hence get eluted after the first passing of the labelled cells. Waste fraction represents cells that are eluted during the column wash prior to the collection of the HSA-selected MDM. The positive fraction represents the HIV-infected HSA-expressing cells retained in the magnetic column that are eluted at the end of the HSA-selection protocol. The gating strategy for detection of HIV-HSA infected macrophages is shown in Fig 8C left panel. The positively selected MDM infected with HIV-HSA showed ∼70% purity while the negatively selected HIV-uninfected cells and waste fractions had ∼7% and ∼10% contaminating HSA expressing MDMs, respectively (Fig 8C, right panel). The results show that RIPK1 was downregulated in the positively selected HIV-HSA enriched fraction compared to the mock infected and negatively selected HIV-uninfected MDM (Fig 8D). **T**hese results indicate that RIPK1 degradation is a consequence of HIV infection of primary MDM.

### SM treatment of HIV-infected MDMs downregulates apoptosis associated signalling molecules TRAF-1 and Bid

In addition to RIPK-1, **t**he process of apoptosis requires the fine-tuned functionality of several signalling molecules including TRAF-1/2, as well as proteins that regulate homeostasis of mitochondria such as Bid and Bax [64,66–68]. We determined the expression of these signalling molecules in response to SM-LCL161 treatment of *in vitro* HIV_CS204_-infected MDM. HIV infection resulted in the down regulation of TRAF-1 (Fig 8E, lanes 1 and 4). Treatment of HIV-infected MDM with SM-LCL161 also resulted in the downregulation of TRAF-1 compared to the mock infected and SM-treated MDMs (Fig 8E lanes 2, 3 and 5). However, TRAF-2 and Bax did not show a significant change in their expression in mock-and HIV-infected MDM as well as between SM-LCL161 treated mock and HIV-infected MDMs (Fig 8E). Bid was downregulated with increasing concentration of SM-LCL161 in the HIV-infected MDM but not in mock-infected MDM (Fig 8E). Overall, these results suggest that SM dysregulates the expression of apoptosis-associated TRAF-1 and Bid in HIV-infected MDM.

### cIAP1/2 and RIPK-1 are essential for survival of MDM

The above results showing inactivation of RIPK1 in settings where cIAPs are absent, may affect the viability of MDM. To determine the combined impact of knockdown of cIAPs and RIPK-1 in the survival of MDM, MDMs from healthy donors were pretreated with necrostatin-1, a specific RIPK-1 inhibitor, for 2 hr followed by treatment with SM-LCL161 and analysis for cell death. Treatment with SM-LCL161 or necrostatin-1 alone did not induce significant cell death of normal primary MDMs. However, combination treatment of SM-LCL161 and necrostatin-1 resulted in significant increase in cell death of primary MDM (Fig 9A). Figure 9B shows representative histograms of the intracellular PI stain. Moreover, treatment with necrostatin-1 alone did not show cleavage of PARP or caspases-8 and 9 although treatment with SM-LCL161 alone did show their minimal cleavage (Fig 9C). However, treatment with both necrostatin-1 and SM-LCL161 revealed significantly enhanced cleavage of the three caspases as well as PARP. **T**hese results suggest that cIAP1/2 and RIPK-1 play an important role in regulating viability of primary human MDM. Since HIV infection down regulates RIPK-1 in MDMs and degradation of IAPs with SM-LCL161 results in death of HIV-infected MDM suggest that RIPK-1 and IAPs play crucial roles in SMs-induced cell death of HIV-infected MDM.

**Figure 9.**
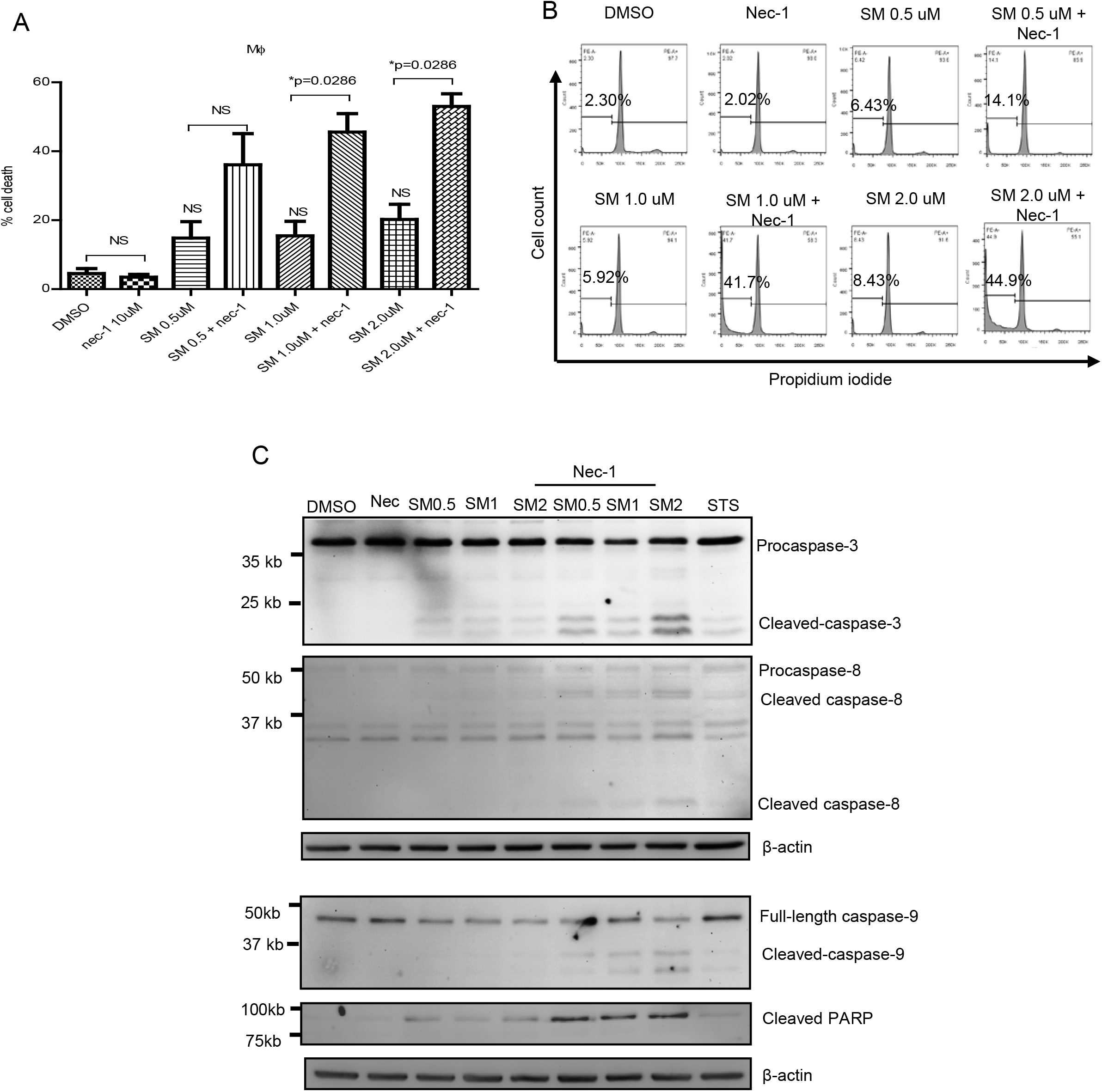
Concomitant downregulation of cIAP1/2 and RIPK-1 in MDMs derived from healthy donors results in activation of apoptosis. (A) MDMs were treated with 10 μM necrostatin-1 for 2 hr followed by the addition of increasing concentration of SM-LCL161 for 48 hr. Cell death was assessed by intracellular PI staining and flow cytometry. p-values were calculated using Mann-Whitney U test (n=4). (B) Representative histograms of the four experiments is shown. (C) MDMs treated as above with necrostatin-1 and SM-LCL161 were harvested and subjected to Western immunoblotting for caspase-3, −8, -and 9, PARP, and beta-actin. The blots shown is a representative of three experiments.

## Discussion

In this study, we investigated the role of IAPs in resistance to apoptosis of HIV-infected Mϕ. We show that although cIAP1/2 are dispensable host factors for the viability of Mϕ, it plays a critical role in the survival of HIV-infected Mϕ. This is illustrated by the observation that SMs induce apoptosis of chronically HIV-infected U1 cell line, *in vitro* HIV-infected MDMs, and *ex-vivo* derived MDMs from naïve and ART-treated HIV patients. SMs were shown to specifically kill HIV-infected MDMs by employing a HSA expressing R5-laboratory strain, HIV-Bal-HSA, and GFP-expressing, HIV-eGFP. The involvement of IAPs was confirmed by employing IAP siRNAs that resulted in killing of HIV-infected MDMs. Our data suggests that SM-induced apoptosis of HIV-infected Mϕ is mediated by apoptosis, is independent of TNF-α and the establishment of M1 polarization. Furthermore, SM-induced apoptosis of HIV-infected Mϕ may be due to RIPK-1 degradation which in concert with IAP1/2 degradation results in apoptosis of HIV-infected Mϕ.

To achieve eradication of HIV-1 in patients undergoing suppressive ART, it is imperative to devise strategies to eliminate HIV reservoirs in cell targets other than T cells such as Mϕ. Recently, IAPs were shown as a potent negative regulator of LTR-dependent HIV-1 transcription and leading to the reversal of HIV latency in JLat latency model system and primary T cells [49]. In addition, IAP1/2 and survivin, another member of the IAP family were suggested to be involved in survival of HIV-infected CD4+ T cells [46,47]. SM activate the non-canonical NF-κB pathway by virtue of RelA:p50 and RelB:p52 transcription factors which bind to the HIV-1 LTR region and results in the induction of virus transcription in latently infected JLat cell lines [49,69]. In addition, XIAP down regulation by flavopiridol, a cyclin-dependent kinase 9 (CDK-9) inhibitor, resulted in increased apoptosis of ACH2 cells (a chronically HIV-infected T cell line) [70]. Additionally, ablation of cIAP1/2 by SMs cleared hepatitis B virus in immune competent mouse models [48]. We and others have previously shown that ablation of cIAP1/2 by SMs does not affect survival of normal primary human Mϕ [51,71]. However, resistance of Mϕ to apoptogenic HIV-Vpr was attributed to cIAP1/2 [51]. These observations suggest that targeting of IAPs may represent a possible strategy for killing of HIV-infected Mϕ. Herein, we show that *in vitro* HIV-infected MDM and MDM generated *ex vivo* from ART-treated or naïve HIV-infected patients were highly susceptibility to SM-mediated cell death. Induction of apoptosis was confirmed by using monomeric (LCL161) and dimeric (AEG40730) SMs as well as IAP-siRNAs and three different M-tropic strains including HIV_cs204_ (clinical), HIV-eGFP and HIV-Bal-HSA. The high variability in the degree of SM-mediated killing of *ex vivo* generated MDM in ART-treated and naïve untreated groups may be due to the effects of antiretroviral drugs on mitochondrial function and highly variable percentage of HIV-infected monocytes in the patients [72,73]. The CD16+ monocytes in the ART-treated patients are significantly higher compared to the untreated group and CD16+ monocytes are more permissive to infection and preferentially harbors HIV-1 *in vivo* [7].

The number of HIV infected Mϕ in *in vitro* infected MDMs is around 5-10% due partly to a milieu of HIV restriction factors that limit the virus life cycle [57,74–76]. However, some of our experiments show killing of around 30% of Mϕ suggesting that SM may cause non-specific killing of bystander MDMs. Since SM did not significantly inhibit virus replication, it is possible that HIV proteins such as Vpr secreted in the supernatants [77] may prime bystander MDMs to SM-mediated killing [50–52]. By employing HIV-Bal-HSA and HIV-GFP strains, our results show that SMs are specifically killing HIV-infected macrophages. Although SM are killing relatively higher number of HIV-HSA-negative MDMs compared to mock-infected MDMs, the differences were not significant. Moreover, SMs killed significantly higher number of HIV-HSA-infected MDMs compared to the HIV-HSA-negative MDMs further suggests the specificity of SMs towards killing of HIV-infected macrophages. cIAP1 was shown to be a negative regulator of LTR-dependent HIV-1 transcription in latently infected primary memory T cells [49]. However, SM did not affect HIV transcription in U1 cells and *in vitro* HIV-infected MDM. We have previously shown that SM treatment alone did not activate either classical or alternative NF-κB pathways in Mϕ [55] that may explain SM’s inability to impact virus replication in Mϕ.

Our results suggest that the mechanism of SM-mediated killing of HIV-infected MDMs is independent of endogenous TNF-α. SM-mediated killing has been attributed to endogenous TNF-α in cancer cells [58,59]; however, it has been reported to be independent of TNF-α in some cancer cells [78]. Given that Mϕ produce high levels of TNFα, the possibility that SM-mediated killing of HIV-1-infected MDM could be attributed to TNF-α was investigated. TNF-α mediated SM-induced killing of myeloid U1 and U937 undifferentiated and PMA-differentiated cells in contrast to that of primary MDMs Although the *in vitro* HIV-infected MDM produced significant amounts of TNFα compared to the mock-infected MDMs, SM treatment did not affect TNF-α secretion in either uninfected or HIV-infected MDM. Moreover, rTNF-α failed to induce cell death in SM-treated MDM suggesting that SM-induced cell death in Mϕ contrary to the cancer cells is independent of TNF-α further displaying dichotomy in the effects of SMs on leukemic myeloid cells and primary macrophages.

HIV infection results in dysregulation of cytokine profile *in vivo* and *in vitro* [79] and can possibly affect the polarization state of Mϕ. Since IFNγ-generated M1 Mϕ are highly susceptible to SM-mediated cell death, the possibility of *in vitro* HIV-1 infection to polarize Mϕ into M1 phenotype making these cells susceptible to SM-induced apoptosis was studied. We show that *in vitro* infected and *ex vivo* derived MDM exposed to SM were not polarized into M1 phenotype suggesting that SM-mediated killing of HIV infected Mϕ was not due to M1 polarization.

Our results suggest that the mechanism of SM-mediated selective killing of U1 cells and primary MDM infected with the clinical strain, HIV_cs204_ is via apoptosis. The pathways of apoptosis are regulated by RIPK-1 [64,67]. In TNFα-mediated signalling, RIPK-1 is recruited in a multiprotein complex I along with TRADD, TRAF2, and cIAP1/2 to promote transcription of genes with anti-apoptotic properties such as cIAP1/2 [67]. RIPK1 is also recruited in a protein complex composed of TRADD, FADD, and caspase-8, which depending on additional proteins recruited, can induce apoptosis or necroptosis [67]. Recently, HIV infection of primary activated CD4^+^ T cells was shown to downregulate RIPK-1 through HIV-1 protease [65]. RIPK-1 modification in response to human rhinovirus and Newcastle disease virus infection has also been reported [80,81]. Herein, we show that infection of MDM with HIV-_CS204_ or with HIV-Bal-HSA caused downregulation and cleavage of RIPK-1. Given that down regulation of IAPs alone by SM LCL161 or of RIPK-1 alone by necrostatin did not induce cell death in uninfected MDM, suggests that RIPK-1 and IAP1/2 are dispensable in survival of Mϕ. However, inactivation of RIPK-1 by necrostatin-1 following IAP degradation by SM resulted in a dramatic increase in cell death, cleavage of caspases and PARP in normal MDM suggesting that RIPK-1 may play a key role in SM-induced killing of HIV-infected Mϕ. The role of RIPK-1 degradation during HIV-1 infection of Mϕ needs further investigation.

TRAF1 is an important receptor interacting protein that forms a complex with TRAF2 to transduce TNFα-induced MAPK and NF-κB activation [82]. TRAF2 is also a key determinant for SM-induced degradation of cIAP1/2 [82]. Our results show that in vitro HIV infection as well as SM-treatment of HIV-infected MDM resulted in downregulation of TRAF-1 but not of TRAF2. In addition, Bid, a proapoptotic protein, is downregulated in SM-treated MDM. Bid is localized in an inactive form in the cytosol which becomes activated by proteolytic cleavage of active caspase-8 [68]. Upon activation, cleaved Bid translocates to mitochondria and forms a complex with Bax to disrupt its integrity resulting in the release of apoptogenic factors, caspase-3 activation and cell death. How SMs cause down regulation of Bid and TRAF-1 in HIV-infected macrophages is not clear. We have shown previously that HIV-Vpr targets Bid, TRAF1 and TRAF2 for proteosomal degradation leading ultimately to mitochondrial outer membrane depolarization and apoptosis [52]. Since SM did not inhibit virus replication and that HIV-Vpr is one of the early genes expressed in virus life cycle, and HIV-Vpr is released following *in vitro* infection of Mϕ, the interplay between Vpr and SM-mediated effect may lead to down regulation of Bid and TRAF-1 and cell death of HIV-infected Mϕ. The functional relevance of the modulation of apoptosis related genes in response to SM-treatment of HIV-infected Mϕ needs further investigation.

In summary, the results of this study suggest the potential significance of SM in killing of HIV-infected Mϕ *in vivo*. In the event SM are able to kill HIV-infected Mϕ *in vivo,* they have the potential to be translated into clinical interventions aimed at eradicating HIV infection by directly targeting HIV-infected Mϕ.

## Materials and methods

### Generation of human monocyte-derived macrophages (MDM), cell lines and reagents

Human peripheral blood mononuclear cells (PBMCs) were isolated by density gradient centrifugation using Ficoll Paque (GE Healthcare Life Sciences Buckingmshire, UK) from the blood of healthy donors. Human MDMs were generated from monocytes via adherence methods as previously described [55]. Briefly, 2.0×10^6^ PBMCs/well were allowed to adhere for 3 hr and non-adherent cells were washed off. Adherent monocytes were cultured for 7 days in complete DMEM (Wisent Inc., St. Bruno, Quebec) medium supplemented with 10% fetal bovine serum (FBS; GE Healthcare), penicillin and streptomycin and 10 ng/mL MCSF (R&D Systems, Minneapolis, MN, USA). MCSF-containing media was replaced every 2 days until the 7^th^ day at which point the monocytes differentiated into macrophages. Purity of macrophages as assessed by measuring CD14 expression by flow cytometry was 100%.

U937 and U1 cells were obtained from NIH AIDS reagent program and were cultured in complete DMEM media. For differentiation, 5.0×10^5^ U937 and U1 cells were treated with 20 nM PMA (Sigma Aldrich, St. Louis, Missouri, USA) for 2 days. The smac mimetics (SM) LCL161 (Active BIochem, Hongkong), and AEG40730 (Tocris Bioscience, Bristol, UK), necrostatin-1 (ApexBio, Houston, TX, USA), staurosporine (ApexBio, Houston, Texas, USA), and LPS (Sigma Aldrich, St. Louis, Missouri, USA) were purchased.

### HIV-1 production and infection of MDMs

The dual tropic HIV-_CS204_ was a gift from Dr. J. Angel (The Ottawa Hospital, Ottawa, ON, Canada). HIV_CS204_ stocks were produced in CD8^+^ depleted PBMCs from healthy donors as described earlier [83]. Stocks of mock virus were produced under similar conditions but in the absence of HIV. HIV growth was determined by measuring p24 using HIV-1 p24^CA^ capture kit as per the manufacturer’ directions (AIDS & Cancer Virus Program, NCI, Fredrick, MD).

The plasmids HIV Gag-iGFP_JRFL (NIH) and pUC-19 (Thermo Fisher Scientific, Waltham, MA, USA) were purchased. The plasmid pNL4.3-Bal-IRES-HSA (provided by Dr M. Tremblay, Laval University, Quebec, Canada) was amplified using One Shot^®^ Stbl3^(TM)^ competent *E. coli* (Invitrogen, Carlsbad, CA, USA) and isolated using endotoxin-free plasmid DNA isolation mega kit (Thermo Fisher Scientific). To produce HIV-1-eGFP, HIV_NL4.3-IRES-Bal_-HSA (HIV-Bal-HSA) and mock viruses, 50 µg endotoxin-free plasmid DNA were transfected into 293T cells with 125 µl of Lipofectamine^TM^ 2000 (Invitrogen) at a density of 18.0 x10^6^ cells/T150 Dish. The supernatants harvested at 48 and 72 hr were combined and centrifuged at 2000 g for 15 min. PEG-it^TM^ virus precipitation solution (SBI, Biotech, Japan) was used to precipitate viruses at 4°C for 24∼48 hr and centrifuged at 2000 g for 30 min. The precipitates were resuspended in PBS with 0.05M HEPES and stored at −80°C. Viruses were quantified by HIV p24 ELISA as above.

The MDMs were infected with 100 ng p24 per 1.0×10^6^ cells supplemented with 5 μg/mL polybrene (Sigma Aldrich) for 16 hr following which cells were infected with HIV_CS204,_ HIV-Bal-HSA or HIV-eGFP for 7 days. The supernatants were assessed for p24 using HIV p24^CA^ ELISA capture kit as above.

### Treatment of MDM with SM or siRNA and assessment of apoptosis

MDMs were cultured in complete media without antibiotics for 2 hr before treatment with various concentrations of SM-AEG40730 or SM-LCL161. For siRNA treatment, 20 nM of siRNA mixture (XIAP, cIAP1 and cIAP2) was added to 200 µl of Opti-MEM^TM^Reduced Serum Medium with 1.0 µl DharmaFect 3 (Dharmacon, Colorado, USA). MDMs were evaluated for cell death by using intracellular PI staining as described [51,84]. Briefly, cells were washed with PBS and fixed with methanol for 15 min at 4 °C. Subsequently, cells were treated with 25 μl of 10 μg/ml RNase A, followed by staining with 25 μl of 1 mg/ml PI solution (Sigma-Aldrich) at 4 °C for 1 h. The DNA content was analyzed using a FACSCanto flow cytometer (BD Biosciences, Franklin Lakes, NJ, USA) and the FACSDiva software. The subdiploid DNA peak (<2N DNA), immediately adjacent to the G_0_/G_1_peak (2N DNA), represents apoptotic cells and was quantified by histogram analyses. PI histograms figures were obtained with WinMDI version 2.8 software (J. Trotter, Scripps Institute, San Diego, CA).

For quantification of apoptosis in HIV-HSA-or HIV-GFP-infected MDMs following treatment with either SM or siRNA transfection, cells were harvested after trypsinization with 0.25% Trypsin-EDTA (Gibco, Dublin, Ireland) for 30 min. HIV-eGFP-infected MDMs were stained with 2.0 µl Annexin-V-APC (BD Biosciences) in 50 µl of PBS/0.5% BSA for 15 min, and fixed with 1% PFA (Affymetrix, Santa Clara, CA, USA). Cells were analyzed with flow cytometer (BD LSR FORTESSA X-20) in GFP and APC channels. HIV-Bal-HSA-infected MDMs were blocked with FcR blocking reagent (MACS Miltenyi Biotec, Auburn, CA, USA) in 50µl of PBS/0.5% BSA, and stained with 2.0 µl of FITC conjugated anti-mouse CD24 antibody (BD Biosciences) for 20 min. Cells were washed and stained with Annexin-V-APC for 15 min, fixed with 1% PFA followed by analysis with flow cytometer at FITC and APC channels.

### Isolation of HIV-HSA-infected MDMs

MDMs were infected with HIV-Bal-HSA for 9 days followed by magnetic sorting using HSA-CD24 beads (MACS Miltenyi Biotec.) through column separation, as previously described [57]. Briefly, infected MDMs were detached with accutase (Innovative Cell Technologies, San Diego, CA), FcRγII receptors were blocked with FcR blocker (MACS Miltenyi Biotec), stained with primary CD24-biotin conjugated antibody and incubated with anti-biotin ultra pure microbeads (MACS Miltenyi Biotec). HSA-expressing cells were collected by positive selection in LS columns. The HSA-negative (negative fraction) cells were collected after passing the labelled cells through the column for the first time. The column was detached from the magnet and the HSA positively labelled cells were collected by plunging out the cells. Purity of the HSA-infected macrophages was assessed by flow cytometry using anti-Biotin PECy7 antibody.

### Analysis of caspase activation

Activation of caspase-3, −8, and −9 was measured as per Abcam’s Caspase staining kit protocol (Abcam, Toronto, Ontario, Canada) by flow cytometry.

### TNF-α ELISA and cytokine ELISA array

Human TNF-α duo set (R&D System) was used to quantify TNF-α as per the manufacturer recommendations. Briefly, the 96-well plates were preincubated with TNF-α capture antibody for 16 hr followed by blocking with 1% FBS. TNFα (1-1000 pg/mL) was used as standards. The samples were added to the plates for 16 hr followed by the detection antibodies for two hr. Next, 100uL/well of substrate solution was added. The enzymatic reaction was stopped with 50 uL/well of stop solution (BioFX Labs, Owing Mills, MD). The plates were read at 490 nM using iMark Microplate reader (Biorad, Mississauga, Ontario) using microplate manager 6 software.

The levels of secreted cytokines were measured as per the directions in Milliplex map kit (Millipore, Etobicoke, ON, Canada). IL-17F, GM-CSF, IFNγ, IL-10, CCL20/MIP3a, IL-12p70, IL-13, IL-15, IL-17a, IL-22, IL-9, IL-1β, IL-33, IL-21, IL-23, IL-5, IL-6, IL-17ε/IL-25, IL-27, IL-31, TNFα, TNFβ, and IL-28A were detected using antibody-immobilized magnetic beads and were quantified by MAGPIX® multiplex with xPONENT® software (Luminex Corp.).

### Western immunoblot analysis

The lysates were subjected to SDS-PAGE electrophoresis as described earlier [51,55,84]. Proteins were transferred onto polyvinylidene difluoride membrane (BioRad Laboratory, Hercules, CA) and probed with primary antibodies specific for cIAP1, cIAP2, XIAP, caspase-3, caspase-8, caspase-9, β-actin, PARP, Bax, TRAF-1, TRAF-2, and RIPK-1 (Cell Signalling Tech, Inc., Danvers, MA), followed by goat anti-rabbit or anti-mouse secondary polyclonal antibodies conjugated to horseradish peroxidase (BioRad Laboratory). Proteins were visualized by enhanced chemiluminescence (Amersham Bioscience, Little Chalfont, UK).

### Statistical analysis

Data was plotted using Graphpad Prism 5. Statistical significance was calculated using student t test or One-way Anova, followed by Dunnett post test. Plotted data represent the mean ± SD.

### Ethics statement

Healthy participants involved in the study gave informed written consent and the protocol for obtaining blood samples was approved by the Review Ethics Board of the Ottawa General Hospital and the Children’s Hospital of Eastern Ontario, Ottawa, ON, Canada.

## Acknowledgements

This study was supported by grants from the Canadian Institute of Health Research to AK (HOP 98830; HOP-107542) and by The Canadian HIV Cure Enterprise Team Grant HIG-133050 (AK) from the CIHR in partnership with CANFAR and IAS. RC was supported by a scholarship from the destination 2020 Faculty of Medicine, University of Ottawa, Ontario, Canada. We acknowledge the generous help provided by the nurses in collecting blood samples. We also acknowledge the healthy donors and HIV patients for providing their blood samples.

The authors declare no conflicting financial interests.

## Authors Contribution

RC, SD and NG performed and analyzed experiments, RC wrote the manuscript, HA performed experiments, EC and JA provided critical insights in writing the manuscripts, JA and WC provided clinical samples, RK provided reagents, MT provided HSA and GFP labelled HIV strains, AK designed the project, analyzed experiments and wrote the manuscript

## Online supporting information

**Supp Fig 1. SM induce the activation of caspases in HIV-infected MDMs.** Human MDMs were *in vitro* infected with HIV-_CS204_ (100 ng p24 / well) for 7 days. The cells were then treated with SM-LCL161 for 48 hr. The activation of the caspase-3, 8, and 9 were detected by intracellular caspases staining and flow cytometry. Representative histograms are shown.

**Supp Fig 2. SMs induce cell death in M1 macrophages.** (A). M0 and M1 MDMs were treated with increasing concentration of SM-LCL161 48 hr (n=3). Cell death was assessed by intracellular PI staining and flow cytometry. The p-values were calculated using Mann-Whitney U test. (B) A representative histograms for cell death in M1 macrophages is shown.

**Suppl. Fig 3. HIV infection of MDMs does not result in the upregulation of cytokines related to M1 phenotype.** MDMs were *in vitro* infected with mock or HIV_CS204_. The supernatants collected after 7 days of infection were analyzed for the secretion of cytokines using Human Th17 magnetic panel cytokine array kit for 22 different cytokines (n=6). The p-values were calculated using Mann-Whitney U test

**Supp Fig 4. SM does not induce aberrant production of M1 cytokines in mock and HIV-infected MDMs. The** *in vitro* mock and HIVcs204-infected MDM for 7 days were treated with SM LCL161 for 48 hr. Supernatants were collected, and cytokine profile was analyzed through Human Th17 magnetic panel cytokine array kit (n=3)

**Supp 5. Supp Fig 6. SM does not induce cytokine production in MDMs generated from HIV-infected individuals. PBMC** from ART-treated HIV+ patients were differentiated into macrophages for 7-days and subsequently treated with SM LCL161 for 48 hr. The supernatants were analyzed for cytokines through Human Th17 magnetic panel cytokine array. P-values were calculated using paired-T test (n=3)

